# Baseline microbiomes of the pillar coral *Dendrogyra cylindrus* reveal novel taxa and regional differences

**DOI:** 10.64898/2026.04.11.717891

**Authors:** Allison Cauvin, Lisa Carne, Kristen L. Marhaver, Mark J.A. Vermeij, Nicolas S. Locatelli, Iliana B. Baums, Valerie J. Paul, Julie L. Meyer

## Abstract

The pillar coral *Dendrogyra cylindrus* is a rare but iconic member of Caribbean reefs that has suffered range-wide losses. *D. cylindrus* is highly susceptible to stony coral tissue loss disease (SCTLD), and the outbreak has contributed to the functional extinction of Florida’s population of pillar corals. The coral microbiome can impact the health and disease resistance of coral colonies, yet little is known about what constitutes the core microbiome of *D. cylindrus*. This information is crucial for comparisons of healthy and diseased tissue in pathogen identification studies and can be applied to restoration efforts as a coral health metric. Therefore, we characterized the microbiomes of *D. cylindrus* colonies ahead of the SCTLD disease front in Belize and Curaçao. The most prevalent members of the *D. cylindrus* microbial community were bacteria for which taxonomy could not be assigned confidently beyond the level of domain as well as the putatively endosymbiotic genera *Endozoicomonas*, *Ca*. Amoebophilus, and *Spiroplasma*. The coral reefs of Belize and Curaçao represent distinct Caribbean marine ecoregions, and we documented regional differences in strains among predominant bacterial taxa. The understudied microbiome of *D. cylindrus* harbors unique bacterial lineages that are in danger of extinction along with its critically endangered coral host, and these bacterial lineages may be important bioindicators during restoration efforts.

**IMPORTANCE:** Tropical corals face global extinction if average temperatures rise by 2°C (3.6°F), which may occur as soon as 2050. Included in the loss of charismatic macrofauna like the majestic pillar coral is the loss of the biological and genetic diversity of its symbionts. Here we examined the bacterial and archaeal communities associated with Caribbean pillar corals and found that the microbiome was dominated by taxonomically unclassified and putatively endosymbiotic taxa. Endosymbiotic bacteria, which live inside the coral tissue, are likely to have evolved unique adaptations to become symbionts and may be important to the health and success of pillar corals in ecosystem restoration efforts.

## Introduction

Pillar corals (*Dendrogyra cylindrus*) are iconic members of Caribbean reefs. Despite being an uncommon or rare species, they are conspicuous due to their towering columnar morphology. As such, they have been referred to as the ‘unicorns’ and ‘magic castles’ of Florida’s reefs. *D. cylindrus* serves as an important source of taxonomic, structural, and biological diversity for Caribbean reefs. It is taxonomically unique as the only member of its genus and behaviorally unique as a coral species that extends its polyps both day and night. *D. cylindrus* also serves as the exclusive host for the dinoflagellate species *Breviolum dendrogyrum* (1, 2), as well as a symbiotic species of shrimp (3). Additionally, *D. cylindrus* serves the role of encouraging aggregation of other marine species. Grunts (reef fishes from the family Haemulidae) aggregate amongst living and dead pillars of *D. cylindrus* (*4*), and the loss of the species could reduce the available three-dimensional structure for vertebrate and invertebrate commensals to take shelter.

Unfortunately, regional populations of pillar corals are under threat. *D. cylindrus* is listed as ‘Critically Endangered’ on the IUCN Red List (5), as well as ‘Endangered’ under the United States Endangered Species Act as of February 2025 (6). Populations of *D. cylindrus* along Florida’s Coral Reef have been in decline in recent decades, primarily due to thermal stress events, a lack of sexual reproduction, and disease outbreaks (1, 7). The emergence of stony coral tissue loss disease (SCTLD) has decimated Florida *D. cylindrus* populations (7, 8). Pillar corals are considered highly susceptible and are among the first in an affected area to exhibit disease lesions (9). Infection of *D. cylindrus* with SCTLD often results in whole colony mortality. Pillar corals are now considered to be nearing total extinction in Florida, with very few genets remaining and distances >10 m separating genetically distinct and reproductively active colonies, decreasing the likelihood of sexual reproduction (7). SCTLD has since spread throughout the Caribbean and negatively impacted regional *D. cylindrus* populations (10, 11). Consequently, this species has been prioritized for conservation and restoration activities, including genetic rescue, *ex situ* propagation, and sample collection and preservation ahead of the SCTLD disease front (7, 12).

When studying an organism in the context of disease, it is important to consider the role of the microbiome. Bacterial symbioses can drive the ecological success and adaptability of their eukaryotic hosts during environmental perturbations (13–15). Coral-associated microbes perform critical functions for the coral host, including nutrient cycling, the production of beneficial compounds such as B vitamins and antibiotics (16, 17), and protection from pathogens (18). For example, coral endosymbionts from the genus *Endozoicomonas* play a vital role in coral host health by producing antimicrobial compounds, competitively excluding pathogenic bacteria, and cycling sulfur via DMSP production (19, 20). In fact, *Endozoicomonas* species show cophylogeny with their coral hosts, reflecting a long-standing symbiotic relationship (21). Coral microbiomes are intimately linked with host phenotype, as it has been demonstrated that coral-associated microbes may contribute to bleaching resilience and resistance to disease (22–24).

Accordingly, understanding drivers of coral microbiome structure and differentiation can have implications for host resilience in the face of compounding disturbances (25, 26). Variation in coral microbial communities can be explained by the coral host species (21, 27), and these communities are distinct from surrounding seawater (21, 28). This implies a degree of microbial selection or filtering imposed by the coral host. While intraspecific coral microbiomes may be influenced by host genet (29, 30) and local environmental conditions (31), broadly they tend to remain stable temporally and geographically (19, 27, 32).

The overall objective of this study was to develop a baseline characterization of the microbiome of pillar coral colonies prior to the arrival of SCTLD. Specifically, this study aimed to identify the core *D. cylindrus* microbiome, or the set of microbial taxa characteristic of a specific host and/or environment (33). Additionally, we sought to examine how these microbial communities may differ across broad geographic ranges as well as the reef-scale variations of putative endosymbionts associated with *D. cylindrus*. The characterization of the *D. cylindrus* microbiome is of interest to the conservation and restoration of this iconic coral, yet little work has been done thus far to characterize the associated microbiome (34). Understanding the baseline microbiome of these corals could facilitate more rapid identification of potential pathogens or disease-causing agents.

## Methods

### Sample Collection

Coral tissue samples were collected from *D. cylindrus* colonies in two countries, Belize (*n*=43) and Curaçao (*n*=128) between November 2019 and May 2020 (collection dates provided in Supplemental Table 1) by collaborators at the Caribbean Research and Management of Biodiversity (CARMABI, Curaçao) and Fragments of Hope (Belize). Most samples were collected between November 2019 and February 2020, a temporal window that represents a relatively consistent thermal regime. Multiple sites within each region were sampled, totaling 9 from Belize and 12 from Curaçao (Figure 1). The three northern sites in Belize (Bacalar Chico Marine Reserve, Tuffy Hol Chan Marine Reserve, and Hol Chan Marine Reserve Table) had reported SCTLD at the time of sampling, though samples were only taken from unaffected colonies. Curaçao did not report its first case of SCTLD until April 2023 (35), well after the coral samples were collected. Coral samples were collected on SCUBA using a hammer and chisel from protruding areas of the colony that were surrounded on all sides by healthy tissue (i.e., no samples were collected from a colony edge that might have been overgrown by CCA or turf algae). Underwater, the samples were stored in separate compartments of a pill organizer filled with seawater. Upon return to shore, these fragments were clipped with bone cutters and placed into 2 ml cryovials filled with 96% ethanol. Between samples, the bone cutters were sterilized by wiping them with ethanol and a Kimwipe. All sample handling was done wearing gloves.

**Figure 1.**
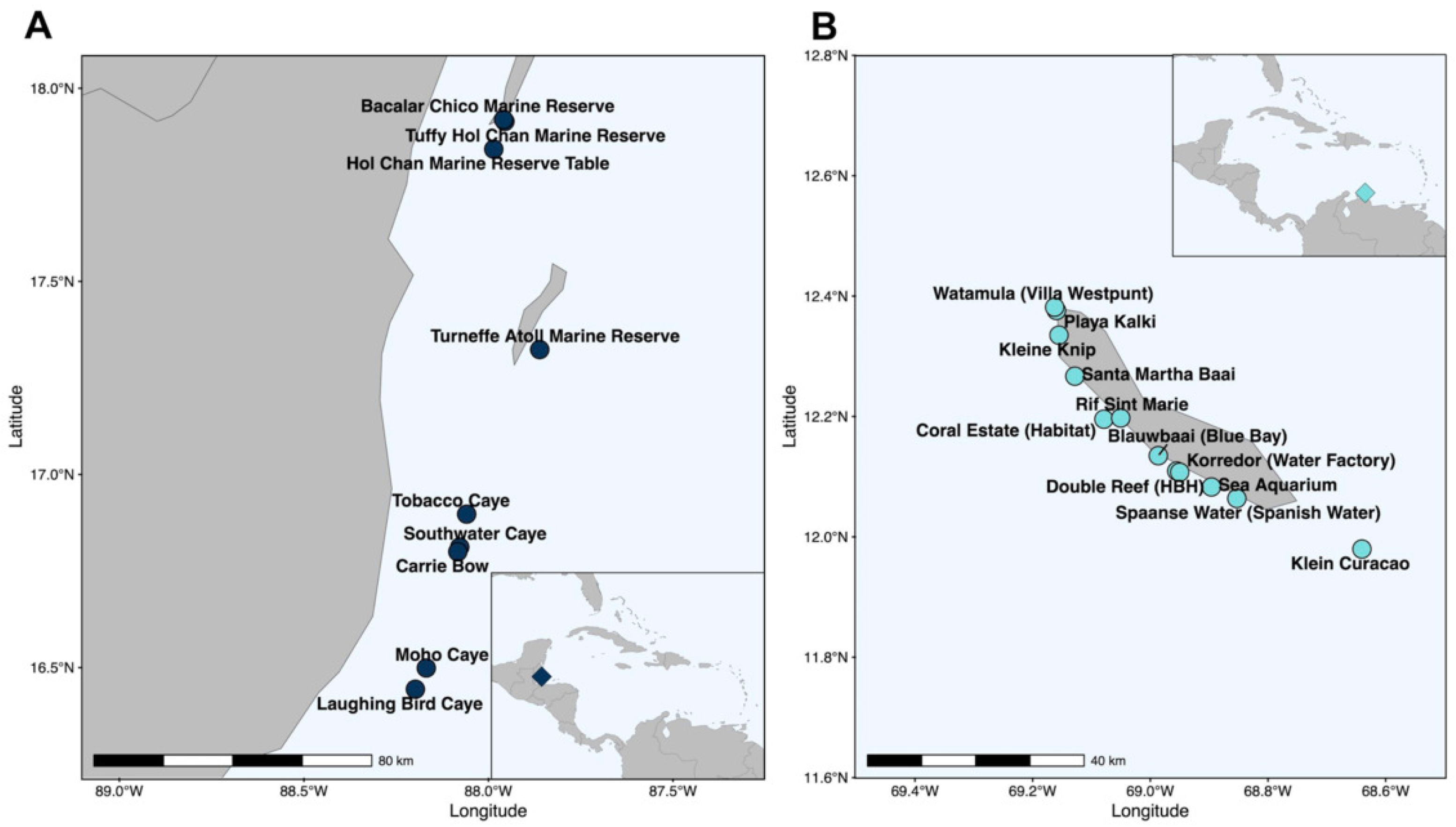
A map of the sampling locations of *D. cylindrus* tissue in Belize (A) and Curaçao (B).

### DNA Extraction & Amplification

DNA was extracted from ethanol-preserved tissue samples using the DNeasy PowerSoil Kit (Qiagen, Hilden, Germany) according to the manufacturer’s protocol. The V4 hypervariable region of the 16S rRNA gene was then amplified via the Earth Microbiome PCR protocol (36) using the primers 515F (5’-GTGGYCAGCMGCCGCGGTAA-3’) (37) and 806R (5’-GGA CTA CNV GGG TWT CTA AT-3’) (38). A single round of PCR was conducted to amplify the V4 region and ligate on the Illumina adapters and barcodes. Each sample was run in triplicate 25 μl reactions. Each reaction included 12.5 μl of Phusion High Fidelity Master Mix (includes Taq; New England Biolabs, Ipswich, MA), 1.25 μl of primer (500 nM final concentration), 0.75 DMSO, and 1 μl of DNA template. All PCR was conducted on an Applied Biosystems SimpliAmp Thermal Cycler (Waltham, MA, USA). Thermocycling conditions were as follows: initial denaturation of 3 mins at 94° C, followed by 35 cycles of denaturation at 94° C, primer annealing at 50° C, and extension for 90s at 72° C, with a final extension of 10 mins at 72° C. Samples were then visualized on an ethidium bromide-stained Tris-acetate-EDTA (TAE) with 1% agarose gel to confirm amplification of the expected product size and that there was no contamination of no-template controls. Amplicon replicates (*n*=3 per sample) were pooled and purified with the Qiagen MinElute PCR Purification Kit according to the manufacturer’s instructions. Purified amplicons were then sequenced on the Illumina MiSeq 2×150bp platform (v2 cycle format) at the University of Florida’s Interdisciplinary Center for Biotechnology Research (RRID:SCR_019152). Each sample was re-sequenced on three separate MiSeq runs to overcome low output of high-quality sequences, which may have been caused by the presence of potential inhibitors in the samples.

### Sequence Processing & Taxonomic Assignment

Sequencing reads were demultiplexed by the sequencing center and primer and adapter sequences removed via cutadapt (39). Sequences from each sequencing run were separately processed with DADA2 (40). The DADA2 bioinformatic pipeline was used for quality assessment, filtering of low-quality reads, error estimation, merging of reads, dereplication, removal of chimeras, and determination of amplicon sequence variants (ASVs). The filtering parameters filterAndTrim(fnFs, filtFs, fnRs, filtRs, truncLen=c(150,150), maxN=0, maxEE=c(2,2), truncQ=2, rm.phix=TRUE, compress=TRUE, multithread=TRUE) were used. The SILVA v138.1 database was used for ASV taxonomy assignment (41, 42). Predominant ASVs that could not be assigned taxonomy beyond the domain or phylum level were compared to the NCBI core nucleotide BLAST database using BLASTn (43). The resulting ASV tables from each sequencing run were then merged using the mergeSequenceTables() function of DADA2. The resulting merged ASV table and sample metadata were then imported into the R package ‘phyloseq’ (44) for further statistical analysis. Reads corresponding to mitochondria, chloroplasts, eukaryotes or those that could not be classified as Bacteria were removed. Samples with more than 500 reads across all three sequencing runs and ASVs with a mean read count of at least 5 across all samples were retained for downstream analyses.

### Microbial Community Structure

Alpha diversity was compared using the estimate_richness function of the phyloseq package to calculate Chao1 and Shannon diversity metrics. To assess microbial beta diversity, the quality-filtered read counts were centered log-ratio transformed with the CoDaSeq package (45). The Aitchison distance was calculated to assess differences in microbial community structure via PERMANOVA using the adonis2 function of the vegan package for R (46) and microbial community variance between regions was visualized using Principal Component Analysis (PCA). Beta diversity dispersion, computed as distance to centroid using the betadisper function of vegan, was compared between the two regions with a Mann Whitney-U test. A multinomial classification method was employed to classify microbial ASVs as *D. cylindrus* microbiome generalists, Curaçao specialists, Belize specialists, or rare microbiome members using the clamtest function (47) of the vegan package.

### Identification of SCTLD Bioindicators

While there was some active SCTLD present on the Belize reefs included in this study at the time of sampling, no samples were taken from diseased colonies. To rule out potential inclusion of visually asymptomatic diseased individuals in the analysis, a custom sequence database was created using the ASVs identified in this project. BLASTn was used to query SCTLD bioindicators identified by Becker et al. (48) against the *D. cylindrus*-associated ASVs. The parameters ‘-outfmt 6 -evalue 10e-6 -ungapped’ were used. Only 100% matches across the 126-bp query were included further.

### Determination of Core Microbiome & Differentially Abundant Taxa

To determine broad taxonomic trends in the *D. cylindrus* microbiome, the fifteen most relatively abundant genera regardless of region were calculated. To identify the most prevalent microbial members, the core microbiome was defined as microbial ASVs that were detected in at least 70% of samples included in the study. This threshold was selected to encompass the microbes that were detected in the majority of *D. cylindrus* microbiomes regardless of geographic region. This was calculated using the phyloseq_filter_prevalence command of the metagMisc R package (49). Differentially abundant taxa between the two regions were determined with DESeq2 (50) using the parametric Wald test. P-values were adjusted using the Benjamini-Hochberg procedure and only taxa with adjusted p-values < 0.01 were considered to be significantly enriched in either region. All figures were generated with the R package ggplot2 (51) except for the clamtest plot, which was generated with base R. The R script to reproduce all analyses is available at https://github.com/meyermicrobiolab/Dendrogyra.

### Phylogenetic Analyses

To further investigate the phylogenetic relationships of microbes associated with *D. cylindrus* within the genera *Endozoicomonas* and *Ca.* Amoebophilus, the sequences of ASVs in this study were compared to sequences from other cnidarian-associated microbes, as analyzed in the meta-analysis conducted by McCauley et al. (52). This is a publicly available dataset of cnidarian-associated microbes for comparison. This analysis was conducted to determine if the microbial phylotypes associated with *D. cylindrus* in this study are phylogenetically similar or distinct from other 16S sequences of these microbial genera identified in other cnidarians from the same region. The database was subset to only include ASVs detected using the 515F-806R primer set in healthy, field-based Cnidarians from the Atlantic-Caribbean. Sequences were aligned using the DECIPHER R-package (53) and trimmed to equal lengths using the subseq function of the Biostrings package (54). ASV sequences were then compared to determine which Cnidarian species share similar ASVs to those detected in *D. cylindrus* microbiomes.

## Results

### Sequencing Results

We successfully sequenced microbial communities from a total of 171 *D. cylindrus* colonies from 9 sites in Belize and 12 sites in Curaçao (Figure 1). After removing samples with less than 500 reads, we identified a total of 7,451 ASVs from 132 coral samples (Table 1; *n* = 35 from Belize, and *n* = 97 from Curaçao), representing 1,054 genera belonging to 544 bacterial families. The average number of reads per coral sample was 6,375 (537–60,352) (Supplementary Table 2). Raw sequencing reads are available in the NCBI Sequence Read Archive under BioProject PRJNA1189597. After filtering for low-abundance ASVs (mean read count of at least 5 across all samples), 183 ASVs remained in the final dataset.

**Table 1.** A summary of the *D. cylindrus* microbiome samples remaining in the analysis after filtering by sequencing depth, broken down by region, site, and sampling dates.

| Region | Site | Sampling Date | Number of Corals |
| --- | --- | --- | --- |
| Belize | Bacalar Chico Marine Reserve | 5/24/20 | 2 |
|  | Tuffy Hol Chan Marine Reserve | 5/22/20 | 1 |
|  | Hol Chan Marine Reserve Table | Date unknown | 2 |
|  | Turneffe Atoll Marine Reserve | 2/25/20 | 4 |
|  | Tobacco Caye | 1/29/20 | 3 |
|  |  | 2/26/20 | 1 |
|  |  | Date unknown | 1 |
|  | Southwater Caye | 2/24/20 | 11 |
|  |  | 3/4/20 | 3 |
|  | Carrie Bow | 3/5/20 | 4 |
|  | Moho Caye | 2/5/20 | 1 |
|  | Laughing Bird Caye | 2/6/20 | 1 |
|  |  | 3/4/20 | 2 |
|  | Site Unknown | 3/5/20 | 2 |
| Curaçao | Klein Curaçao | 11/13/19 | 8 |
|  | Spaanse Water (Spanish Water) | 2/5/20 | 8 |
|  | Sea Aquarium | 2/3/20 | 18 |
|  | Double Reef (HBH) | 1/21/20 | 10 |
|  | Korredor (Water Factory) | 1/19/20 | 3 |
|  | Blauwbaai (Blue Bay) | 2/5/20 | 6 |
|  | Coral Estate (Habitat) | 2/24/20 | 1 |
|  | Rif Sint Marie | 2/24/20 | 15 |
|  | Santa Martha Baai | 2/3/20 | 9 |
|  | Kleine Knip | 2/3/20 | 4 |
|  | Playa Kalki | 1/20/20 | 3 |
|  | Watamula (Villa Westpunt) | 2/24/20 | 12 |

### Alpha and Beta Diversity Metrics

Microbial communities from Belize and Curaçao corals did not differ in alpha diversity by either Chao1 or Shannon diversity metrics (p > 0.05). Microbial community composition differed by region, though the effect size was small (PERMANOVA, R^2^ = 0.060, p = 0.001) (Figure 2A). Sites within regions also explained variation in the structure of *D. cylindrus*-associated microbial communities in Curaçao (R^2^ = 0.201, p = 0.001), but not in Belize (R^2^ = 0.241, p=0.702). Beta diversity dispersion, as measured by distance to centroid, was higher in *D. cylindrus* microbiomes from Curaçao compared to those from Belize (p = 0.0004) (Figure 2B). Using a multinomial species classification method, 15.8% of the ASVs were categorized as *D. cylindrus* microbiome generalists, 67.8% as Curaçao specialists, and 16.4% as Belize specialists (Figure 3, Supplementary Table 3). No ASVs in this dataset were categorized as too rare for classification.

**Figure 2.**
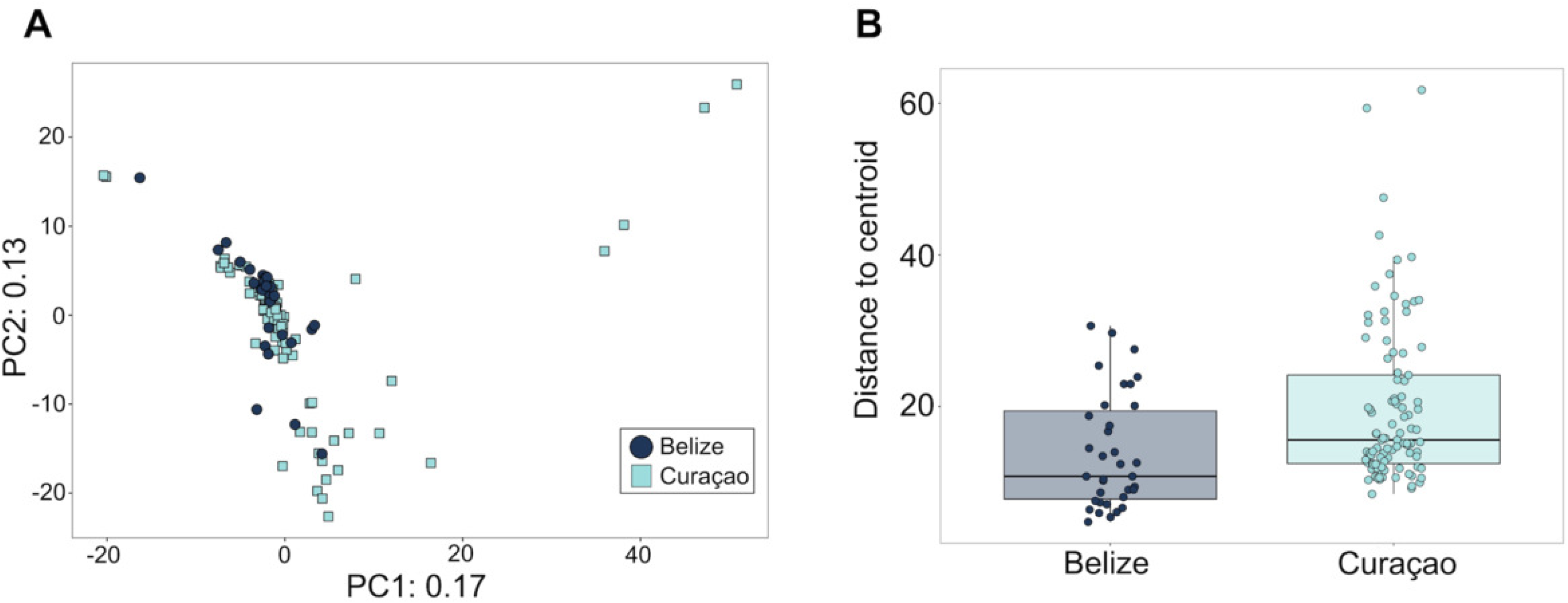
**A)** Principal component analysis (PCA) of *D. cylindrus* microbial community structure in Belize and Curaçao. Each axis represents variance within the microbial community structure. **B)** Beta diversity dispersion computed as distance to centroid of *D. cylindrus* microbiomes between Belize and Curaçao.

**Figure 3.**
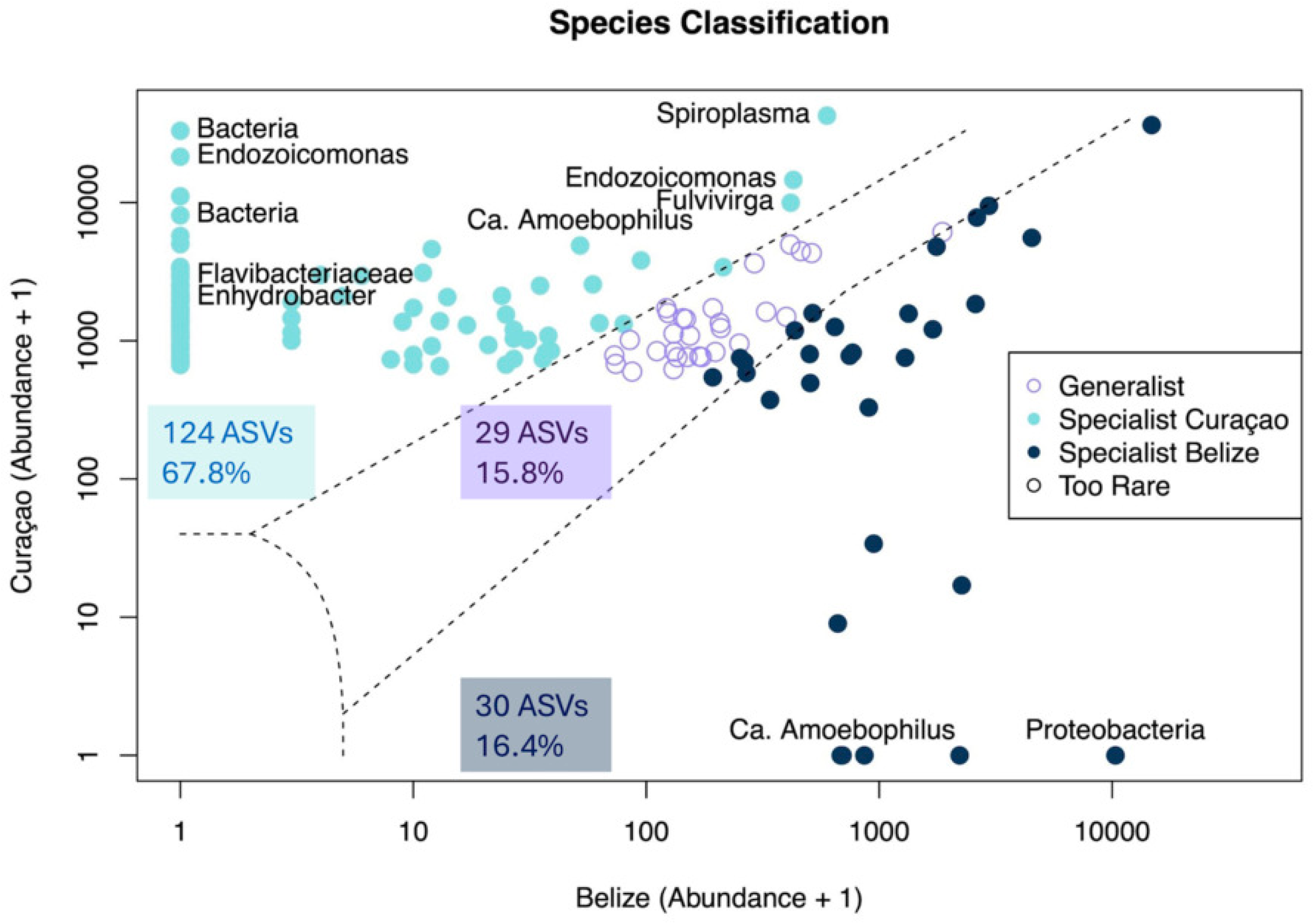
Amplicon sequence variants (ASVs) were classified either as *D. cylindrus* generalists (purple), Curaçao specialists (light blue), or Belize specialists (dark blue) based on a multinomial species classification method (clamtest). There were no ASVs considered too rare to classify. The 11 ASVs detected as differentially abundant between Curaçao and Belize are labeled in the plot.

### SCTLD Bioindicators Were Identified but Not Abundant

The BLASTn query of SCTLD bioindicators against the *D. cylindrus*-associated ASVs yielded 13 exact sequence matches (Supplemental_file_1.fasta) in the unfiltered dataset of 7,451 ASVs. These included five *Vibrio* ASVs, two *Fusibacter* ASVs, as well as *Shimia*, *Thalassobius*, Arcobacteraceae, *Halarcobacter*, *Algicola*, and *Halodesulfovibrio* ASVs. Only one *Vibrio* SCTLD bioindicator ASV (ASV38) was detected in the filtered dataset of 138 ASVs, demonstrating that most of the SCTLD-bioindicators were of very low relative abundance. The *Vibrio* ASV38 was detected in 24 out of 132 samples with an average relative abundance of 4.9% in those 24 samples.

### Abundant Microbial Genera, the Core Microbiome, and Differentially Abundant ASVs

The 15 most relatively abundant bacterial genera in the *D. cylindrus* microbiome included putatively endosymbiotic genera such as *Endozoicomonas* (Phylum *Pseudomonadota*), *Ca*. Amoebophilus (Phylum *Bacteroidota*), and *Spiroplasma* (Phylum *Bacillota*), as well as other genera commonly associated with coral, including *Vibrio* and *Ruegeria* (Figure 4). In addition, a substantial portion of the *D. cylindrus* microbiome is composed of bacteria with unresolved taxonomy, including three ASVs that could only be classified to the domain Bacteria and one ASV that could only be classified to the phylum *Proteobacteria* (now *Pseudomonadota*) with the SILVA 138.1 database. Thus, we further compared these ASVs to the NCBI core nucleotide BLAST database. Two of the three unclassified bacterial ASVs were 97 - 98% similar to the 16S rRNA gene in a metagenome-assembled genome of a *Spirochaetia* strain (Phylum *Spirochaetota*) from a Florida algae metagenome sequenced through the Aquatic Symbiosis Genomics Project (NCBI GenBank Accession OZ253665.1). The third unclassified bacterial ASV as well as the unclassified proteobacterial ASV were 97% and 95% similar, respectively, to a clone library sequence (NCBI GenBank Accession JQ236445.1) from the Mediterranean coral *Balanophyllia europaea*, which is a member of the same coral family (*Dendrophylliidae*) as *D. cylindrus*. The unclassified proteobacterial ASV was exclusively detected in Belize samples. Notably, the most abundant ASV in an unpublished microbiological characterization of Florida *D. cylindrus* from *in situ* and *ex situ* coral nurseries (34) was also an unclassified bacterial ASV whose closest match was the same clone sequence (JQ236445.1) from the Mediterranean *Dendrophylliidae* coral.

**Figure 4.**
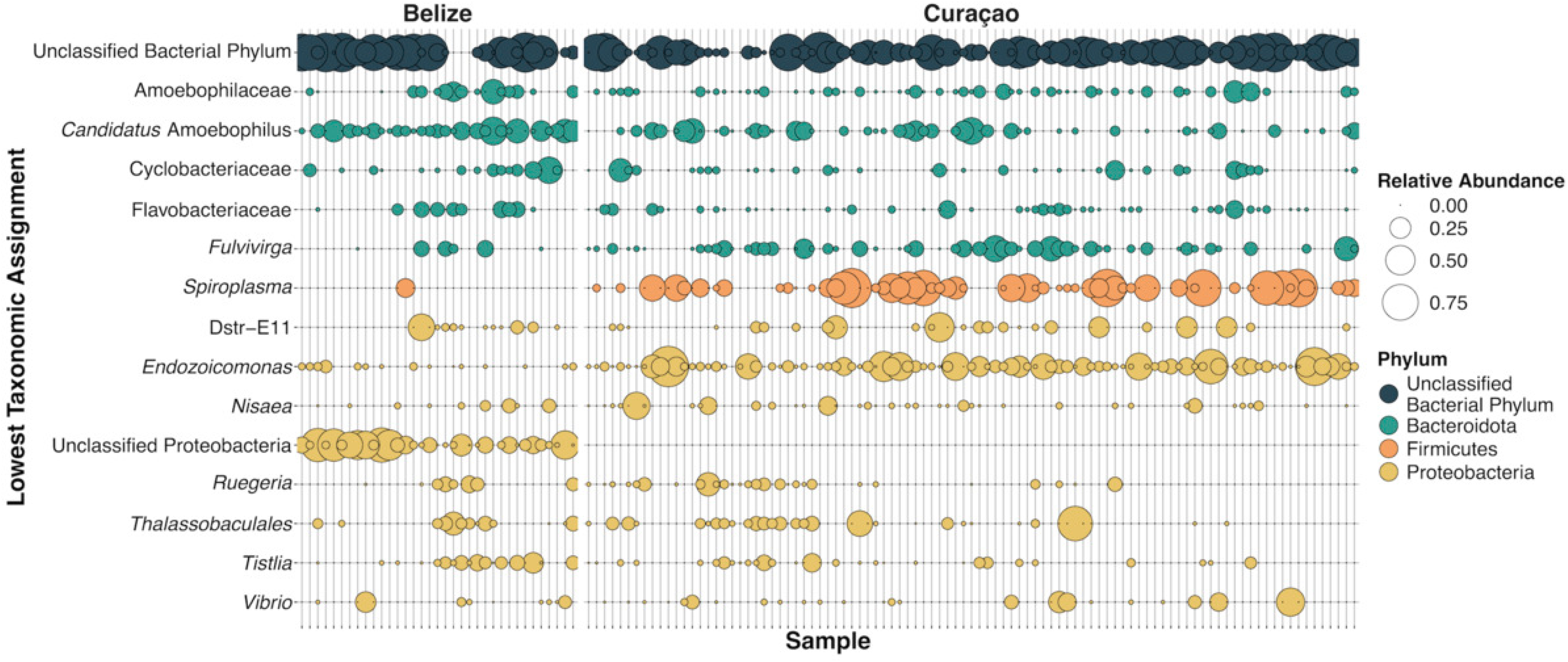
The 15 most abundant genera in *D. cylindrus*, labeled with the lowest taxonomic assignment. Each point represents the relative abundance of that genus within each sample.

Only two ASVs exceeded the 70% prevalence threshold to be considered core members of the *D. cylindrus* microbiome: one unclassified ASV whose closest match was the *Spirochaetia* metagenome-assembled genome (OZ253665.1) and one *Endozoicomonas* ASV. The *Endozoicomonas* ASV, detected in both Belize and Curaçao, had 100% nucleotide identity to a long list of clone library sequences from *Porites lutea, Porites astreoides,* and *Orbicella annularis* (for example, NCBI GenBank Accessions KF180058.1, GU119042.1, DQ200445.1).

DESeq2 analysis identified 11 ASVS that were differentially abundant between Belize and Curaçao (Figure 5). One unclassified Proteobacteria ASV and one *Ca.* Amoebophilus ASV were more abundant in Belize, while ASVs of unclassified Bacteria, *Endozoicomonas*, *Enhydrobacter*, unclassified Flavobacteriaceae, and *Fulvivirga* were more abundant in Curaçao.

**Figure 5.**
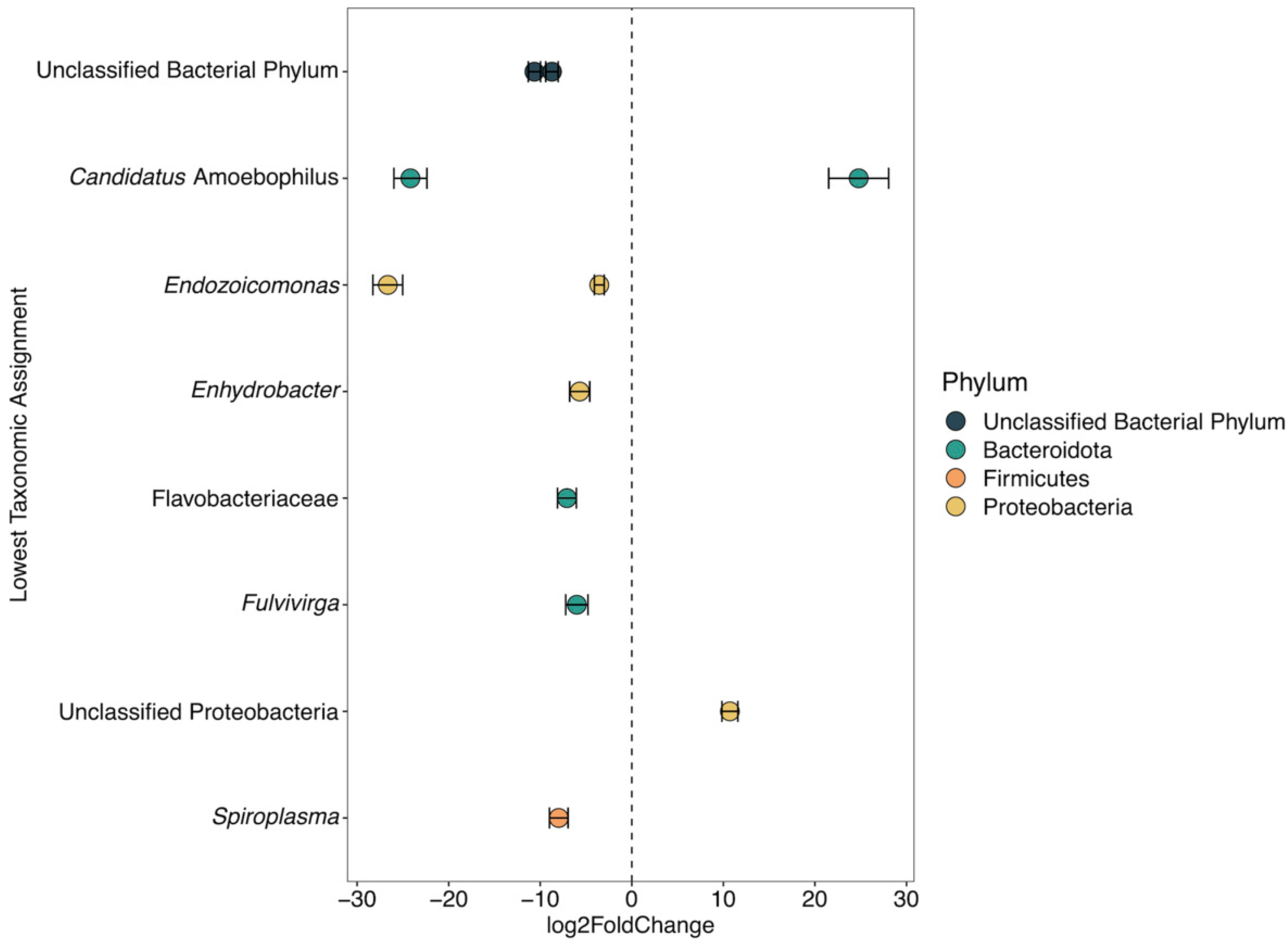
The log2fold change of the 11 ASVs detected as differentially abundant between Belize and Curaçao. Taxa on the left side of the dotted line were enriched in Curaçao (negative change) and taxa on the right side of the dotted line were enriched in Belize (positive change) by DESeq2.

### Putative Endosymbionts in the D. cylindrus Microbiome

Notably, three putatively endosymbiotic genera were found to be among the most abundant in *D. cylindrus* microbiomes and exhibited regional differences in prevalence and abundance: *Endozoicomonas*, *Ca*. Amoebophilus, and *Spiroplasma*.

Seven *Endozoicomonas* ASVs were detected in *D. cylindrus* microbiomes, though only two (*Endozoicomonas* ASV1 and ASV2) are predominant on Curaçao and Belize reefs (Figure 6A). *D. cylindrus* microbiomes from Curaçao harbored more abundant and more diverse *Endozoicomonas*, as all 7 ASVs were detected on Curaçao reefs. In contrast, only *Endozoicomonas* ASV2 was detected in *D. cylindrus* microbial communities in Belize. In the McCauley dataset (52), 51 *Endozoicomonas* ASVs were detected in Atlantic-Caribbean cnidarian microbiomes with the same primers used in this study and included in comparative analyses. After aligning these sequences and trimming them to equal lengths, the McCauley dataset contained 42 unique *Endozoicomonas* ASVs, while six unique *Endozoicomonas* ASVs were from the *D. cylindrus* corals in this study (*Endozoicomonas* ASV3 and ASV4 from *D. cylindrus* detected in this study were identical after sequence trimming to match the McCauley dataset sequence length). Three ASVs from the *D. cylindrus* corals in this study were detected in other Atlantic-Caribbean cnidarian microbiomes (Figure 6C, Table 2), including ASV2 that was detected on both Belize and Curaçao reefs and was identified as part of the core microbiome. Additionally, three *Endozoicomonas* ASVs found in *D. cylindrus* corals from this study were not found in cnidarians from the meta-analysis.

**Figure 6.**
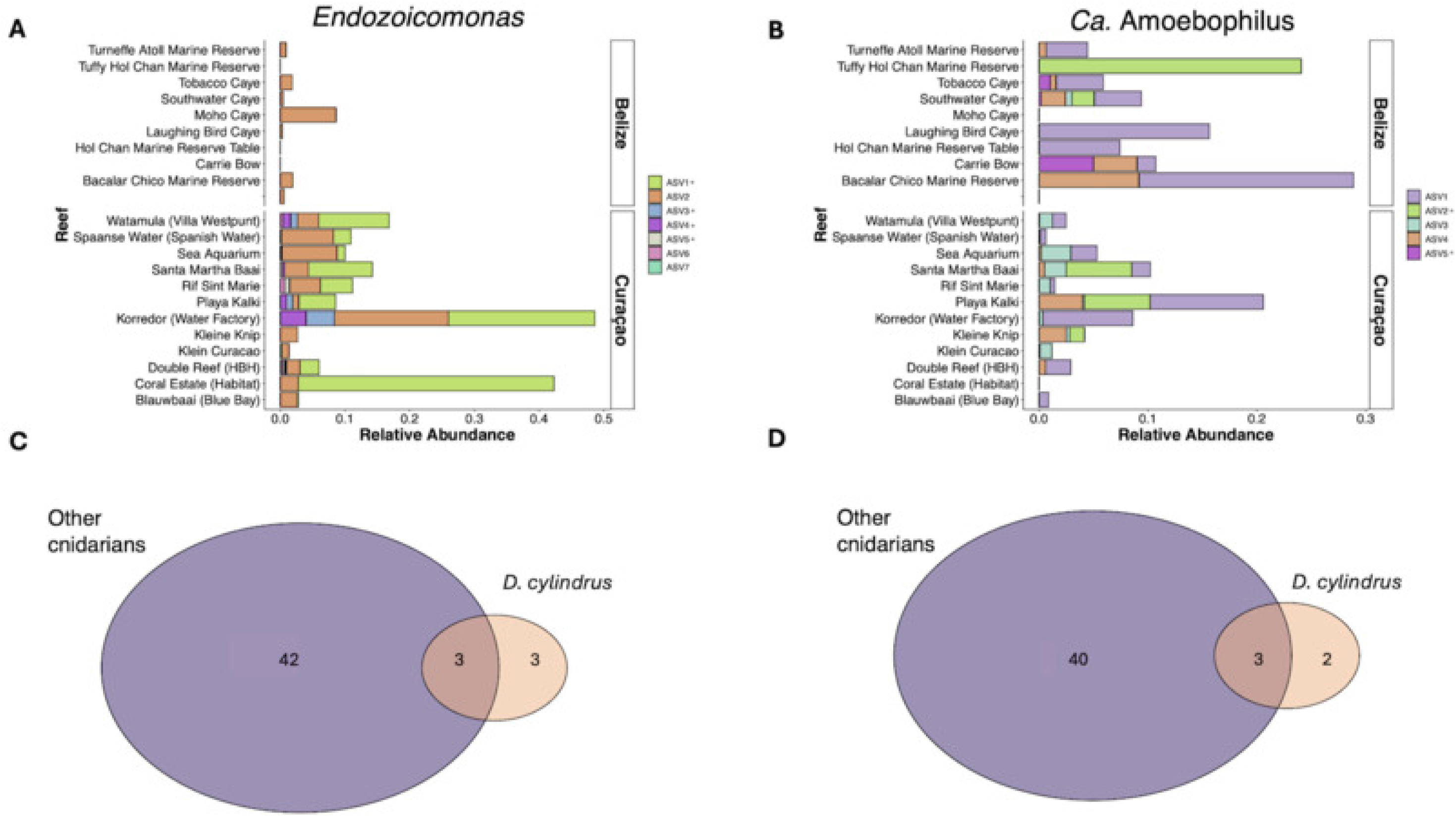
A summary of the presence of two predominant bacterial genera in *D. cylindrus* microbiomes. **A)** The average relative abundance of each *Endozoicomonas* ASV on each reef. Each bar represents the abundance of *Endozoicomonas* on reefs in Belize and Curaçao, colored by each ASV. ASVs designated by * are unique to *Dendrogyra cylindrus*. **B)** The average relative abundance of each *Ca.* Amoebophilus ASV on each reef. Each bar represents the abundance of *Ca.* Amoebophilus on reefs in Belize and Curaçao, colored by each ASV. ASVs designated by * are unique to *Dendrogyra cylindrus*. C) The number of *Endozoicomonas* ASVs shared or distinct between *D. cylindrus* microbiomes and other Caribbean cnidarians. The purple circle shows the number of *Endozoicomonas* ASVs detected in other cnidarians, and the pink circle shows the number of *Endozoicomonas* ASVs detected in this study. **D)** The number *Ca.* Amoebophilus ASVs shared or distinct between *D. cylindrus* microbiomes and other Caribbean cnidarians. The purple circle shows the number of *Ca.* Amoebophilus ASVs detected in other cnidarians, and the pink circle shows the number of *Ca.* Amoebophilus ASVs detected in this study

**Table 2.**
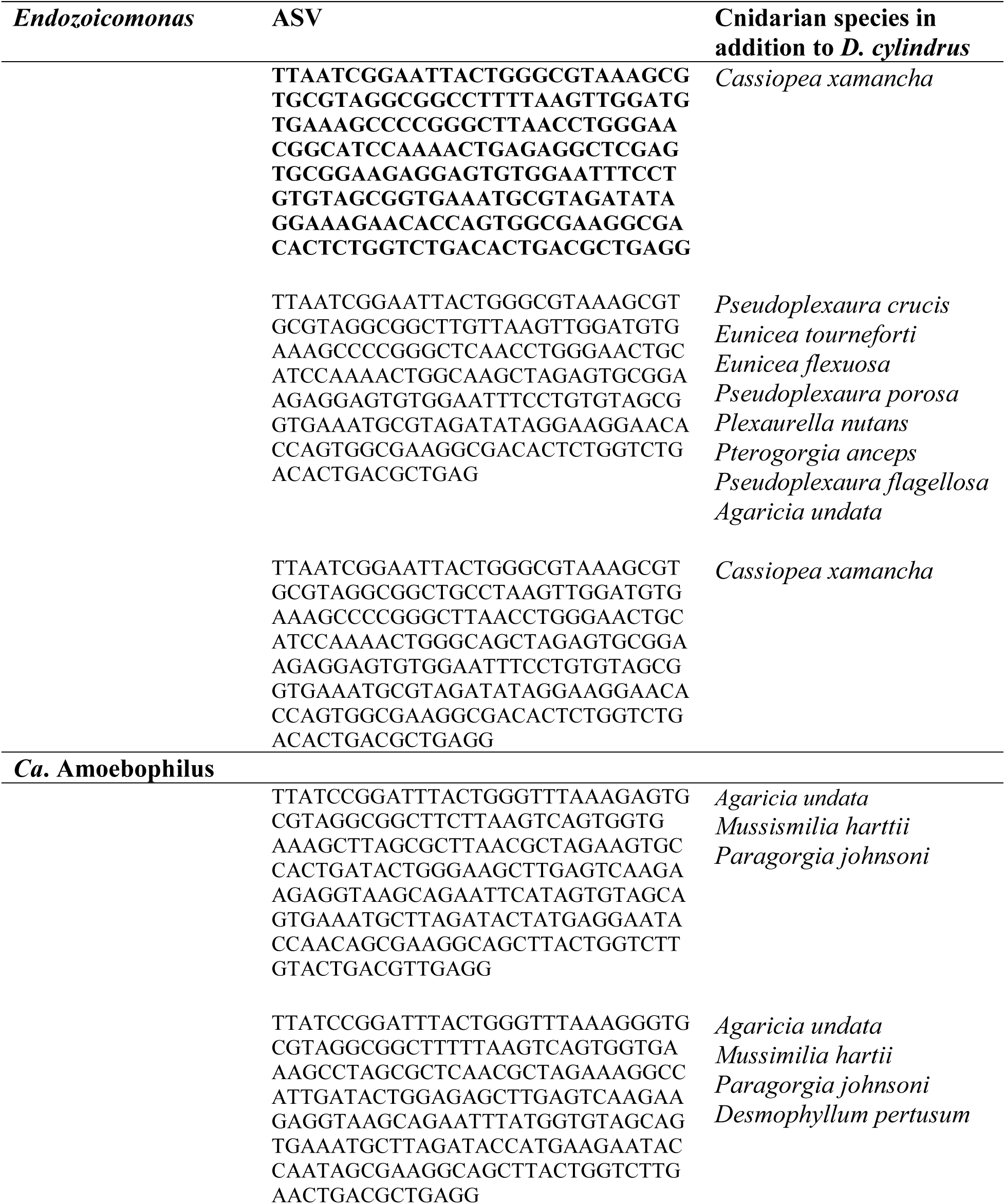
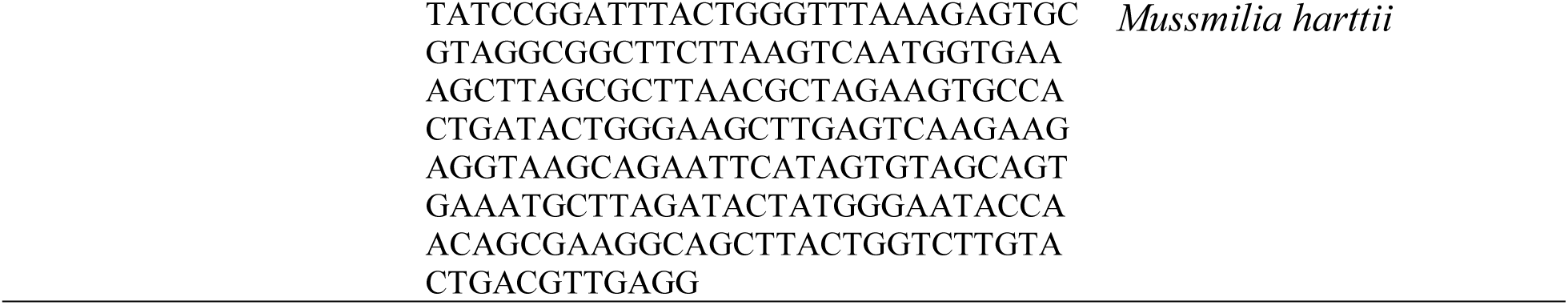
Amplicon sequence variants (ASVs) classified as *Endozoicomonas* and *Ca.* Amoebophilus that are shared between *Dendrogyra cylindrus* and other Caribbean cnidarian species. The bolded *Endozoicomonas* ASV was identified as part of the core microbiome of *D. cylindrus* as it was present at least 70% of samples and detected from both Belize and Curaçao.

Five distinct ASVs of *Ca*. Amoebophilus were also detected in *D. cylindrus* microbiomes, and these ASVs were less geographically distinct when compared to *Endozoicomonas* (Figure 6B). Four of the five *Ca*. Amoebophilus ASVs were detected in both Belize and Curaçao, while only one (ASV5) was found exclusively in Belize. The majority of reefs sampled in this dataset displayed multiple *Ca*. Amoebophilus ASVs. When compared to the McCauley dataset (52), 43 unique *Ca.* Amoebophilus ASVs were detected in Atlantic-Caribbean cnidarian microbiomes. Three of the *Ca*. Amoebophilus ASVs found in this study were shared with other Atlantic-Caribbean cnidarians (Table 2), while two were considered unique to *D. cylindrus* (Figure 6D).

One *Spiroplasma* ASV was detected in 54 out of the 82 Curaçao samples and in one *D. cylindrus* colony from Belize. When compared against the closest NCBI BLAST matches, the *Spiroplasma* ASV was only 90.1% similar to the closest hits, which include *Spiroplasma* spp. from flies (for example, NCBI GenBank Accession PV254006.1). Three *Spiroplasma* ASVs were detected in the McCauley dataset (52), though only two of these were amplified from the same 16S rRNA primer region as the current study and included for further analysis. The two ASVs from the McCauley dataset had 100% similarity to each other over their shared nucleotides but were 92.9% similar to the *Spiroplasma* ASV identified in this study.

## Discussion

*Dendrogyra cylindrus* harbors a unique microbiome that is predominantly composed of putatively endosymbiotic bacteria and bacteria of uncertain taxonomic classification.

*Endozoicomonas*, *Ca.* Amoebophilus, and *Spiroplasma* were abundant and prevalent in *D. cylindrus* microbiomes and members of each of these genera have been detected as endosymbiotic or intracellular, as described below. Additionally, several predominant ASVs in this study could not be classified beyond the phylum or class level. This highlights the limitations of current reference databases for the classification of previously undescribed microbial sequences, as well as the novelty of the *D. cylindrus* microbiome. There is an increased need to improve our reference databases to fully leverage the sequence data generated. More importantly, the presence of multiple distinctive bacteria in this phenotypically and taxonomically unique coral species that harbors a unique algal symbiont means that the extinction of this iconic holobiont would be a tragic loss of novel biodiversity on multiple levels.

We observed regional differences in *D. cylindrus* microbial community structure among the most commonly detected taxa. Some studies have demonstrated that coral host species is an important driver of microbial community structure and found high degrees of similarity in species-specific coral microbiomes across broad geographic ranges (19, 27), indicating that coral hosts can curate their microbiome in a species-specific manner. The variation in microbial communities between Belize and Curaçao suggests that these corals may respond to local environmental conditions by forming associations with specific bacteria that provide some benefit to the holobiont, as proposed by (13). Notably, Belize and Curaçao fall into distinct Caribbean marine ecoregions with different thermal regimes and storm occurrence, supporting the hypothesis that the regional microbiome variations observed here may be environmental in origin (55, 56). We found that the majority of the *D. cylindrus* microbiome (84.2% of ASVs) is composed of regional specialists. Even though our sampling efforts were unequal, resulting in three times as many samples from Curaçao as from Belize, we did not find significant differences in regional alpha diversity. Thus, the increased sampling effort in Curaçao did not lead to a more diverse microbiome. In addition, three of the most abundant taxa in the dataset, including an unclassified bacterial ASV, an *Endozoicomonas* ASV, and a *Ca.* Amoebophilus ASV were detected only in Belize, the location with much fewer samples.

*D. cylindrus* harbors diverse *Endozoicomonas* symbionts, as previously demonstrated in other coral species (57). *Endozoicomonas* is a common member of coral microbiomes globally, especially in coral tissues (58), and is often considered to be a beneficial symbiont of corals (59), associating with multiple coral species in high abundance (20, 60, 61) and demonstrating cophylogeny with their coral host (21). Fluorescent *in situ* hybridization microscopy and laser-capture microdissection have demonstrated the presence of *Endozoicomonas* aggregates located within coral tissues, often in close proximity to *Symbiodinaceae* (62–65). Similarly, intracellular microcolonies of *Endozoicomonas* spp. have been detected in fish and a wide variety of mollusks (66, 67) and some strains can exhibit rapid growth within the host nucleus that results in cell rupture (68). Comparative analysis of diverse marine *Endozoicomonas* genomes corroborates that members of this genus exhibit the capacity for a variety of lifestyles ranging from free-living to obligate symbioses that can be mutualistic to parasitic (69). Members of this bacterial genus possess a variety of genomic mechanisms to maintain a beneficial relationship with their coral host and their algal symbionts, including T3SS secretory proteins that allow these symbionts to regulate gluconeogenesis, scavenge H_2_O_2_, and participate in carbon and sulfur cycling (70, 71), thus allowing them to participate in energy production, nutrient cycling, and holobiont protection. The high prevalence and abundance of two *Endozoicomonas* ASVs in *D. cylindrus* microbiomes may indicate that these particular phylotypes possess metabolic capabilities that benefit their coral host and are therefore more highly selected associates, while others are more transient based on local environmental conditions (72). However, the role of *Endozoicomonas* in coral health remains enigmatic as its presence has been associated with a decreased ability to respond to environmental change (73). In addition, there may be a tradeoff between coral growth and disease susceptibility related to the abundance of *Endozoicomonas* in the microbiome (58). Together, this suggests that the lifestyles of the *Endozoicomonas* strains associated with *D. cylindrus* warrant further study to determine both their physical locality in coral tissues as well as their genetic capacity to either benefit or harm the coral host.

*Ca.* Amoebophilus and other ASVs belonging to the Amoebophiliaceae family were among the most abundant members of the *D. cylindrus* microbiome. *Ca.* Amoebophilus appears to be a ubiquitous microbial member of Caribbean coral tissues, as sequences belonging to this bacterial group have been found in other studies (21, 60, 74) and have also been associated with the shallow layers of the coral skeleton (75). This bacterial group was initially identified as an endosymbiont of the *Acanthoamoeba* genus of amoebas (76), and genome comparison studies have demonstrated that the *Ca.* Amoebophilus genome contains a multitude of protein domains associated with host cell interactions (77), providing a genomic basis for an intracellular lifestyle in this bacterial lineage. While it cannot be ruled out that this bacterial group is residing within amoebas associated with coral rather than the coral tissue itself, its prevalence in cnidarian microbiome metaanalyses (52, 60) and the genomic potential for symbiosis warrant further investigations of this bacterial group and its impact on cnidarian holobiont fitness.

One novel *Spiroplasma* ASV was identified by this study in high prevalence and abundance within the Curaçao samples. While *Spiroplasma* has been detected in *Porites* microbiomes in Mo’orea (23) and in the microbiomes of several Caribbean coral species (52, 78), it is notable as a relatively rare member of Caribbean coral microbiomes. Other bacterial members of the *Spiroplasma* genus have been characterized as widespread vertically transmitted endosymbionts among a variety of arthropod hosts (79). It cannot be ruled out that the *Spiroplasma* ASV sequenced in this study was sourced from marine arthropods associated with *D. cylindrus* rather than inhabitants of coral tissue. If that is the case, the fact that this bacterial group was nearly exclusively detected in Curaçao *D. cylindrus* microbiomes could indicate that these corals are under higher arthropod challenge in Curaçao or that arthropod vectors are differentially infected with *Spiroplasma* between the two regions. Some of the closest matches on NCBI for the *Spiroplasma* ASV included the *Spiroplasma* strain SNeo from the *Drosophila neotestacea* fly, which protects its host from parasitism via the production of toxins (80, 81). *Spiroplasma* endosymbionts can elicit phenotypic changes within their host organism as a result of horizontally acquired genes that encode for toxins or protective genetic elements (80, 82–85). Further genomic analyses such as the generation of metagenome-assembled genomes from coral-associated *Spiroplasma* strains will enable insights into the genetic potential of these microbes to elicit phenotypic changes within their host.

The *D. cylindrus* microbiome contains several distinct putatively endosymbiotic bacterial lineages. It has been proposed that a high abundance of *Endozoicomonas* promotes coral growth through metabolic mutualism but could result in a tradeoff for higher disease susceptibility (69). This is because host physiological characteristics that promote bacterial symbioses could also leave the coral vulnerable to invading pathogens (86) and symbiotic bacteria can engage in immune suppression to maintain evolutionary symbioses (87–89). *D. cylindrus* is particularly vulnerable to several coral diseases, including white plague, black band disease, and SCTLD (7). Further investigation into how microbes modulate host disease susceptibility is warranted, particularly endosymbiotic species which possess a high potential to impact the host phenotype.

The baseline microbial communities characterized in this study can be applied to ongoing *D. cylindrus* conservation and restoration efforts. Recent *ex situ* spawning and propagation efforts have been highly successful (12, 90). In Florida, the rescue of genetically diverse *D. cylindrus* colonies prior to local arrival of SCTLD was conducted through the Association of Zoos and Aquariums Florida Reef Tract Rescue Project (91) and provides diverse genets for *ex situ* sexual reproduction and propagation. An important consideration in these propagation efforts must be the role of the coral microbiome as a biological indicator of coral health (92). The characterization of baseline healthy microbiomes of *D. cylindrus in situ* can be useful for understanding the impact of stressors in *ex situ* rearing environments. Additionally, the results of this work can be applied to understanding the shifts in microbial communities after diseases like SCTLD have impacted the region and what that may indicate for coral health and resilience.

## Acknowledgements

Laboratory work was supported by startup funds from the University of Florida (JM). AC was supported by the University of Florida Graduate School Preeminence Award. NSL was supported by the CBIOS (NIH T32 Kirschstein-NRSA: Computation, Bioinformatics, and Statistics) training program at The Pennsylvania State University (#T32GM102057). Funding for fieldwork and molecular work was provided by the Paul G. Allen Family Foundation (Grant #16004-1 from Multiplier). Samples from Belize were collected and exported under Marine Scientific Research permit 0005-20 from the Belize Fisheries Department and CITES permit 08978. Samples from Curaçao were exported under CITES permit 16US784243/9. This is contribution # XXXX of the Smithsonian Caribbean Coral Reef Ecosystem and contribution # XXXX of the Smithsonian Marine Station.

---

**Supplementary File 1**. Nucleotide sequences of the 13 ASVs that were identical matches to SCTLD bioindicators.

**Supplementary Table 1.** Collection dates and locations for 171 Dendrogyra cylindrus samples.

**Supplementary Table 2.** Number of sequencing reads for 16S rRNA gene library over 3 separate sequencing runs. ASV tables were generated separately for each sequencing run and read counts for each step of the dada2 pipeline are shown.

**Supplementary Table 3.** Amplicon sequence variants (n=183) used in the primary analyses, with classes assigned by the multinomial species classification method and taxonomic identifications.

